# Menace to the ultimate antimicrobials among common *Enterobacteriaceae* clinical isolates in part of North-East India

**DOI:** 10.1101/610923

**Authors:** Mohan Sharma, Pankaj Chetia, Minakshi Puzari, Nakul Neog, Amrit Borah

## Abstract

**Introduction:** *Enterobacteriaceae*, the normal dwellers in the human intestine, commonly associated with a variety of community acquired and nosocomial infections. Emerging trend of antibiotic resistance among these strains is a notable issue globally; more serious threat is the resistance against the available last resort antibiotics- the carbapenems. Our study thus intended to determine the burden of resistance towards this ultimate antimicrobial class, so as to assist in the empiric therapeutic decision making process and to search for alternate options.

**Materials and Methods:** Our study was a cross-sectional study with inclusion of clinical isolates collected from varied sources, from health settings in upper Assam. The isolates were identified based on standard methods of morphology study and biochemical tests. The identified isolates were then subjected to antibiotic susceptibility testing following Kirby-Bauer disc diffusion method and the result interpreted as per the CLSI guidelines. The resistance of the reported carbapenem resistant isolates was confirmed by minimum inhibitory concentration (MIC) determination using commercial E-strip kit.

**Results:** Among the enterobacterial isolates *Klebsiella* spp. accounted the majority, followed by *Escherichia coli*, *Citrobacter* spp., *Shigella* spp. and others. Multi-drug resistance (MDR) was noted among 67.6% isolates; however, carbapenem resistance was confirmed in 18.9% of the total *Enterobacteriaceae* isolates.

**Conclusion:** Higher prevalence of resistance towards the last resort antimicrobial, carbapenems, among the *Enterbacteriaceae* isolates of upper Assam seems to be upcoming threat to the region, limiting the treatment options in future.

## Introduction

*Enterobacteriaceae* is a family of Gram-negative bacteria, normal inhabitant of the human gut, which may spread outside and cause different serious infections such as Pneumonia, Blood stream infection, Urinary tract infection (UTI), Wound infection and Meningitis [1, 2]. Resistance towards different classes of existing antibiotics, among the members of *Enterobacteriaceae*, is an alarming issue worldwide [3]. Carbapenems are a powerful group of broad spectrum beta-lactam antibiotics which, in many cases are our last effective defense against multi-drug resistant (MDR) and extended spectrum beta-lactamase (ESBL) bacterial infections [4, 5]. Increased use and misuse of most of the beta-lactam antibiotics and carbapenems, resulted in the emergence of Carbapenem-resistant *Enterobacteriaceae* (CRE) with worldwide presence [2, 6, 7].

CRE have been associated with high mortality and morbidity rates of up to 40-50% reported in some studies [8]. Several studies on CRE had been conducted in India from time to time; however, there are limited reports on the prevalence and distribution of CRE from the North-East region. A study from Manipur reported 30% of the Gram-negative isolates obtained from a tertiary care hospital were resistant towards carbapenem [9].

Our study thus aimed to bridge the gap of the prevalence data on carbapenem resistance among the *Enterobacteriaceae* isolates obtained from different health settings of upper Assam.

## Methods

The study was a cross-sectional laboratory based study, conducted from March 2017 to September 2018, involving a total of 312 non duplicate enterobacterial isolates obtained from varied samples. The isolates were obtained from the routine clinical samples of selected health settings in four districts of upper Assam, namely, Dhemaji, Dibrugarh, Sivasagar and Tinsukia.

### Screening of *Enterobacteriaceae* isolates and their identity confirmation

Identification of the obtained Gram negative isolates was performed by their colony morphology, Gram staining characteristics, catalase test, oxidase test, and other relevant biochemical tests as per standard laboratory identification methods.

### Antibiotic susceptibility testing

Antibiotic susceptibility testing was done by Kirby-Bauer disc diffusion method on Mueller Hinton agar (MHA) against antibiotics of different classes: Amikacin (AK), Amoxicillin/Clavulanic acid (AMC), Piperacillin/Tazobactum (PIT), Cefotaxime (CTX), Ceftazidime (CAZ), Ceftriaxone (CTR), Cefepime (CPM), Tetracycline (TE), Co-Trimoxazole (COT), Ciprofloxacin (CIP), Levofloxacin (LE), Nitrofurantoin (NIT) and Imipenem (IPM). Zone of inhibition interpreted as per the Clinical and Laboratory Standards Institute (CLSI) guidelines [10]. *Escherichia coli* ATCC 25922 was used as a susceptible control strain. Those strains which were non susceptible to at least one agent in 3 or more antimicrobial categories were reported as MDR and those non susceptible (intermediate or resistant) to a carbapenem were CRE [8].

### Screening of MDR Enterobacteriaceae isolates for confirmation of ESBL production

All the isolates resistant to at least one of the three indicator cephalosporins, namely ceftazidime (30μg), cefotaxime (30μg) and ceftriaxone (30μg) were considered for ESBL production test, using phenotypic ESBL E-strip test (HiMedia, Mumbai)

### CRE detection and confirmation

CRE strains obtained from Kirby-Bauer disk diffusion method were subjected to carbapenem MIC determination using E-strip test (HiMedia, Mumbai)

## Results

### Clinical Sources and the isolate profile

During the course of the study, total of 312 *Enterobacteriaceae* isolates were obtained from various clinical sources; majority of the specimen were urine (62%), followed by sputum (15%), blood (7%), wound swab (6%), pus (3%), throat swab (2%), stool (2%) and other sources (3%) (Fig 1).

**Fig1.**
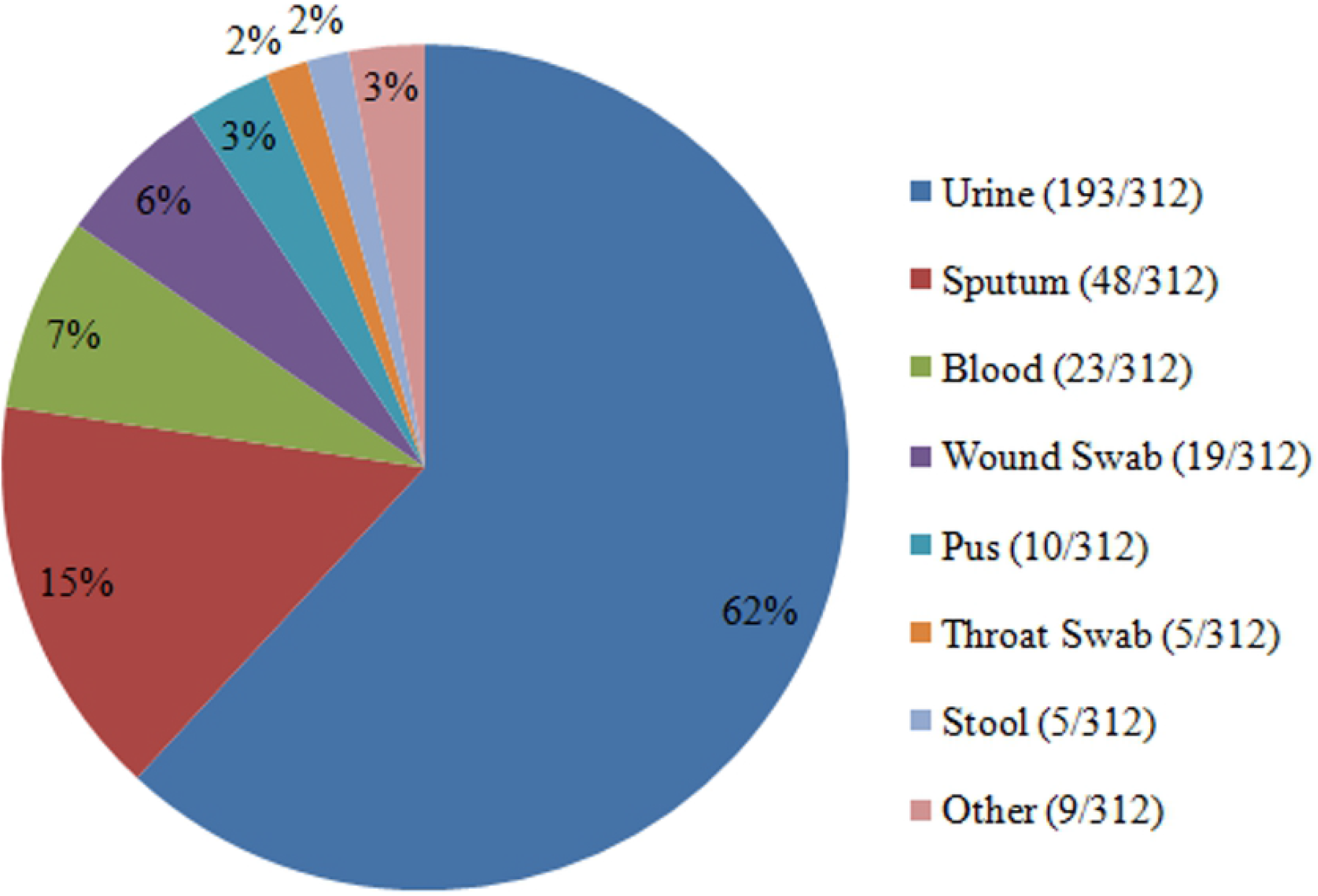
Distribution of the isolates obtained from various clinical sources. *Klebsiella* spp. and *Escherichia coli* accounted the majority 51.3% and 42.0% respectively. Other enterobacteria were less frequent; *Citrobacter* spp. (2.2%), *Shigella* spp. (1.6%), *Enterobacter* spp. (1.3%), *Proteus* spp.(1.0%) and *Yersinia* spp. (0.6%) (Table 1).

**Table1.** Distribution of the enterobacteria isolates against specimen types

| Enterobacterial isolates | Specimen type |  |  |  |  |  |  |  |  |  |  |  |  |  |  |
| --- | --- | --- | --- | --- | --- | --- | --- | --- | --- | --- | --- | --- | --- | --- | --- |
|  | Urine | Sputum | Blood | Wound Swab | Pus | Throat Swab | Stool | Tracheal aspirate | Tracheotomy tube | Aural Swab | CSF | Vaginal Swab | Catheter tip | Peritoneal fluid | Total |
| *Klebsiella spp.* | 71<br>(36.8%) | 41<br>(85.4%) | 19<br>(82.6%) | 14<br>(73.7%) | 4<br>(40.0%) | 5<br>(100%) | 0 | 1<br>(50.0%) | 1<br>(100%) | 1<br>(100%) | 1<br>(100%) | 1<br>(50.0%) | 1<br>(100.0%) | 0 | **160** |
| *E. coli* | 113<br>(58.5%) | 7<br>(14.6%) | 1<br>(4.4%) | 2<br>(10.5%) | 5<br>(50.0%) | 0 | 1<br>(20.0%) | 0 | 0 | 0 | 0 | 1<br>(50%) | 0 | 1<br>(100%) | **131** |
| *Citrobacter spp.* | 3<br>(1.6%) | 0 | 3<br>(13.0%) | 0 | 0 | 0 | 0 | 1<br>(50.0%) | 0 | 0 | 0 | 0 | 0 | 0 | **7** |
| *Enterobacter spp.* | 4<br>(2.1%) | 0 | 0 | 0 | 0 | 0 | 0 | 0 | 0 | 0 | 0 | 0 | 0 | 0 | **4** |
| *Proteus spp.* | 0 | 0 | 0 | 2<br>(10.5%) | 1<br>(10.0%) | 0 | 0 | 0 | 0 | 0 | 0 | 0 | 0 | 0 | **3** |
| *Shigella spp.* | 1<br>(0.5%) | 0 | 0 | 1<br>(5.3%) | 0 | 0 | 3<br>(60.0%) | 0 | 0 | 0 | 0 | 0 | 0 | 0 | **5** |
| *Yersinia spp.* | 1<br>(0.5%) | 0 | 0 | 0 | 0 | 0 | 1<br>(20.0%) | 0 | 0 | 0 | 0 | 0 | 0 | 0 | **2** |
| **Total** | **193** | **48** | **23** | **19** | **10** | **5** | **5** | **2** | **1** | **1** | **1** | **2** | **1** | **1** | **312** |

### Antibiotic susceptibility pattern

The susceptibility pattern of different *Enterobacteriaceae* isolates, against a spectrum of 13 selected antibiotics of different categories, was analyzed (Table 2). There was variable susceptibility towards different antibiotics, non-susceptibility or resistance was highest against amoxicillin/clavulanic acid (65.4%), followed by third generation cephalosporins- ceftriaxone (62.2%), ceftazidime (60.6%) and cefotaxime (51.6%). Susceptibility of piperacillin/tazobactum combination and fourth generation cephalosporin-cefepime was comparatively better, with resistance of 36.2% and 38.8% respectively. Of the total isolates, 47.1% were resistant towards tetracycline. Quinolones exhibited variable resistance, ciprofloxacin and levofloxacin resistance were 46.2% and 37.2% respectively. High percentage of resistance to co-trimoxazole (44.9%) and nitrofurantoin (45.5%) also observed, challenging the use of sulfonamide and nitrofuran class of drugs. Aminoglycoside representative, amikacin was found to be most effective against all isolates with better susceptibility rate with the minimal resistance of 21.8% only. Resistance toward the carbapenem drug- imipenem was 32.4%, though lesser but a threat.

**Table2.** Antibiotic resistance pattern of different *Enterobacteriaceae* isolates

|  | AK | AMC | PIT | CTX | CAZ | CTR | CPM | TE | COT | CIP | LE | NIT | IPM |
| --- | --- | --- | --- | --- | --- | --- | --- | --- | --- | --- | --- | --- | --- |
| *Klebsiella*<br>spp. (n=160) | 40<br>(25%) | 110<br>(68.8%) | 52<br>(32.5%) | 78<br>(48.8%) | 99<br>(61.9%) | 102<br>(63.8%) | 57<br>(35.6%) | 64<br>(40.0%) | 62 (38.8%) | 65<br>(40.6%) | 45<br>(28.1%) | 84<br>(52.5%) | 47<br>(29.4%) |
| *Escherichia*<br>coli (n=131) | 25<br>(19.1%) | 79<br>(60.3%) | 52<br>(39.7%) | 75<br>(57.2%) | 80<br>(61.1%) | 81<br>(61.8%) | 60<br>(45.8%) | 72<br>(55.0%) | 68 (51.9%) | 71<br>(54.2%) | 67<br>(51.1%) | 52<br>(39.7%) | 53<br>(40.5%) |
| *Citrobacter*<br>spp. (n=7) | 0<br>(0.0%) | 3 (42.9%) | 1<br>(14.3%) | 0<br>(0.0%) | 4<br>(57.1%) | 5<br>(71.4%) | 1<br>(14.3%) | 5<br>(71.4%) | 4 (57.1%) | 2<br>(28.6%) | 1<br>(14.3%) | 0<br>(0.0%) | 0<br>(0.0%) |
| *Shigella* spp.<br>(n=5) | 1<br>(20.0%) | 5<br>(100%) | 1<br>(20.0%) | 1<br>(20.0%) | 2<br>(40.0%) | 1<br>(20.0%) | 0<br>(0.0%) | 2<br>(40.0%) | 2 (40.0%) | 2<br>(40.0%) | 1<br>(20.0%) | 1<br>(20.0%) | 0<br>(0.0%) |
| *Enterobacter*<br>spp. (n=4) | 0<br>(0.0%) | 4<br>(100%) | 4<br>(100%) | 2<br>(50.0%) | 1<br>(25.0%) | 1<br>(25.0%) | 2<br>(50.0%) | 0<br>(0.0%) | 2 (50.0%) | 2<br>(50.0%) | 1<br>(25.0%) | 2<br>(50.0%) | 1 (25.0%) |
| ***Proteus* spp.**<br>(n=3) | 2<br>(66.7%) | 3<br>(100%) | 3<br>(100%) | 3<br>(100%) | 3<br>(100%) | 3<br>(100%) | 1<br>(33.3%) | 3<br>(100%) | 1 (33.3%) | 1<br>(33.3%) | 1<br>(33.3%) | 1<br>(33.3%) | 0<br>(0.0%) |
| ***Yersinia* spp.**<br>(n=2) | 0<br>(0.0%) | 0<br>(0.0%) | 0<br>(0.0%) | 2<br>(100%) | 0<br>(0.0%) | 1<br>(50.0%) | 0<br>(0.0%) | 1<br>(50.0%) | 1 (50.0%) | 1<br>(50.0%) | 0<br>(0.0%) | 2<br>(100%) | 0<br>(0.0%) |
| **Total**<br>(N=312) | **68**<br>(21.8%) | **204**<br>(65.4%) | **113**<br>(36.2%) | **161**<br>(51.6%) | **189**<br>(60.6%) | **194**<br>(62.2%) | **121**<br>(38.8%) | **147**<br>(47.1%) | **140**<br>(44.9%) | **144**<br>(46.2%) | **116**<br>(37.2%) | **142**<br>(45.5%) | **101**<br>(32.4%) |

**AK-** Amikacin (30mcg/disc), **AMC-** Amoxicillin/ Clavulanic acid (20/10 mcg/disc), **PIT-** Piperacillin/Tazobactam (100/10 mcg/disc), **CTX-**Cefotaxime (30mcg/disc), **CAZ-** Ceftazidime (30mcg/disc), **CTR-** Ceftriaxone (30mcg/disc), **CPM-** Cefepime (30mcg/disc), **TE-** Tetracycline (30mcg/disc), **COT-** Co-Trimoxazole (1.25/23.75mcg/disc), **CIP-** Ciprofloxacin (5mcg/disc), **LE-** Levofloxacin (5mcg/disc), **NIT-** Nitrofurantoin (300mcg/disc), **IPM-**Imipenem (10mcg/disc).

### Prevalence of Multi-Drug Resistant (MDR) *Enterobacteriaceae*

Of the total isolates considered for the study, 67.6% of the isolates were reported to be MDR. Among major isolates, MDR prevalence was found to be higher within *E.coli* (67.9%), followed by *Klebsiella* spp. (66.9%). Remaining isolates exhibited varied frequency of MDR prevalence according to their numbers concerned (Table 3).

**Table3.**
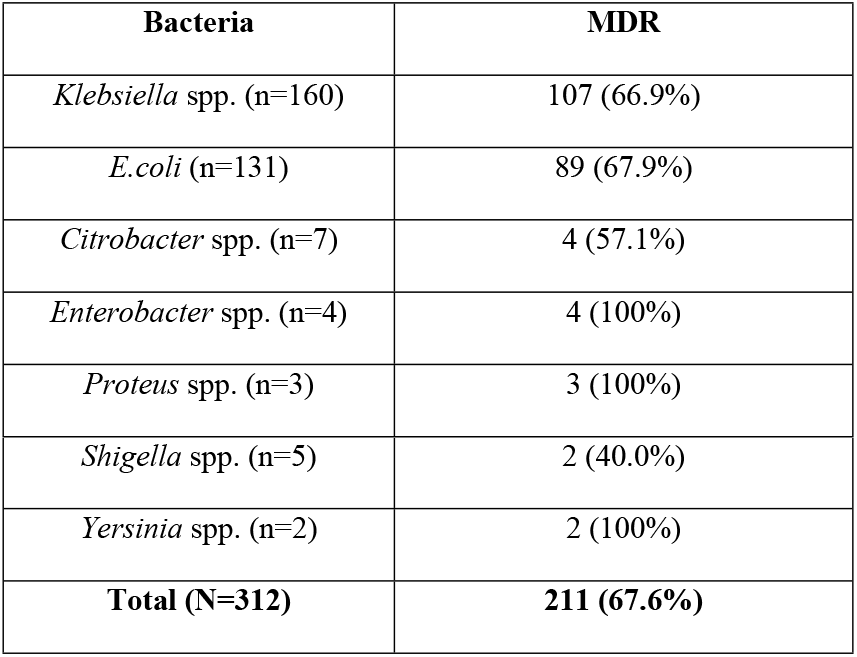
MDR prevalence among different *Enterobacteriaceae* isolates

### Confirmed extended spectrum beta-lactamase (ESBL) producers

Out of 226 isolates, which were screened for ESBL production, overall ESBL producers were 138 (44.2%) as depicted from the result of MIC ratio method, using Ezy MIC strip. ESBL producing trend was found to be the highest among *E. coli*, with 60 isolates (45.8%) were positive. Among *Klebsiella* spp. isolates 72(45.0%) were confirmed to be ESBL producer. Remaining organisms were found to be showing varying pattern of ESBL production (Table 4).

**Table4.**
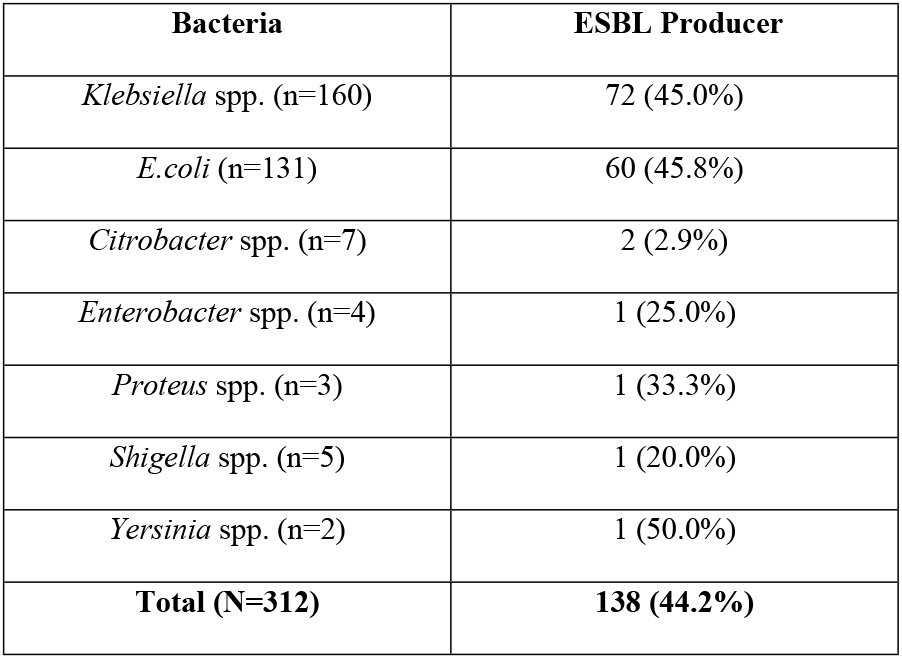
Prevalence of ESBL producers among different isolates based on E-strip test

| **Bacteria** | **ESBL Producer** |
| --- | --- |
| *Klebsiella* spp. (n=160) | 72 (45.0%) |
| *E.coli* (n=131) | 60 (45.8%) |
| *Citrobacter* spp. (n=7) | 2 (2.9%) |
| *Enterobacter* spp. (n=4) | 1 (25.0%) |
| *Proteus* spp. (n=3) | 1 (33.3%) |
| *Shigella* spp. (n=5) | 1 (20.0%) |
| *Yersinia* spp. (n=2) | 1 (50.0%) |
| **Total (N=312)** | **138 (44.2%)** |

### Prevalence of CRE based on MIC

A total of 101 *Enterobacteriaceae* isolates were found to be non-susceptible towards imipenem by Kirby-Bauer disk diffusion method, these included 53 (40.5%) *E.coli* isolates, 47 (29.4%) *Klebsiella* spp. and 1 (25.0%) *Enterobacter* spp. Further confirmation of carbapenem resistance was done by MIC determination, which revealed 59 isolates to confirmed CRE, accounting 18.9% of the total *Enterobacteriaceae* isolates. Interestingly, like the ESBL producers, here also CRE were the highest in *E.coli*, 37 (28.2%) compared to *Klebsiella* spp. 22 (13.8%) isolates. Among other isolates none of them were CRE based on their MIC determination (Table 5).

**Table5.** Prevalence of Carbapenem resistance among the enterobacteria isolates

| **Bacteria** | **Bacteria resistant by Kirby-Bauer disk diffusion method** | **Resistance based on MIC by Ezy-MIC gradient strip** |
| --- | --- | --- |
| *Klebsiella* spp. (n=160) | 47 (29.4%) | 22 (13.8%) |
| *E.coli* (n=131) | 53(40.5%) | 37 (28.2%) |
| *Citrobacter* spp. (n=7) | 0 (0.0%) | 0 (0.0%) |
| *Enterobacter* spp. (n=4) | 1 (25.0%) | 0 (0.0%) |
| *Proteus* spp. (n=3) | 0 (0.0%) | 0 (0.0%) |
| *Shigella* spp. (n=5) | 0 (0.0%) | 0 (0.0%) |
| *Yersinia* spp. (n=2) | 0 (0.0%) | 0 (0.0%) |
| **Total (N=312)** | **101 (32.4%)** | **59 (18.9%)** |

## Discussion

In the whole period of study, urine samples were the major specimen source of the isolates (62.0 %), as reported by other studies. [9, 11] This trend might be due to the fact that urinary tract infection (UTI) is the common problem among the patients visiting the health settings. *Klebsiella* spp. and *E. coli* were reported as the most common *Enterobacteriaceae* isolates, similar to earlier findings [11, 12, 13]. However, the distribution is not in concordance to those observed earlier, in our case *Klebsiella* spp. accounted the highest 51.3% and *E. coli* were 42.0%, which might be due to the variation in the type of the source specimen.

Antibiotic susceptibility pattern shown by the isolates, against the 13 different antibiotics, was variable. Amikacin and Imipenem were found to be effective drugs, with susceptibility rate of 78.2% and 67.6% respectively. Susceptibility towards other drugs considered under the study was mostly lesser than 50.0%, with comparatively higher resistance among the beta-lactams. This pattern is similar to other studies, which might be due the increasing trend of, chromosome or plasmid mediated, β-lactamase production among the enterobacterial strains [14, 15, 16, 17].

Overall prevalence of multidrug resistance (MDR) was reported to be 67.9%, which is a very serious issue and is comparable to the earlier studies conducted in Ethiopia (68.3%) and Nepal (about 65.0%) [18, 19, 20]. Among the *E.coli* isolates 67.9% were reported to be MDR, higher than earlier reports [21, 22]. Within the *Klebsiella* spp. isolates, 66.9% were MDR, which is far less compared to that observed before [23, 24]. These variations might be due to the varied specimen sources and the numbers of the isolates considered in the study.

Among all the *Enterobacteriaceae* isolates 44.2% were confirmed to be ESBL producers, which is near about similar to that observed from Bhopal (48.3%) [12], but higher than that reported by other studies done in different part of India [25, 26]. Also the prevalence rate was observed to be higher compared to the earlier study reported from the same geographic region (24.6%) [27], ESBL prevalence within *E.coli* isolates also found to be 45.8%, higher compared to a similar study carried out in the region [28], but lesser compared to the study of Sharma et al. who reported 56.9% from Jaipur. Similar pattern of *Klebsiella* spp. ESBL producer also noticed accounting to 45.0%, which is somewhat similar to that found in the study from parts of India and Iran [11, 29], but were comparatively lesser than reported from Rajasthan, 67.0% [30]. Our finding of higher ESBL producers among *E.coli*, is in conformity with some of the previous studies [11,13].This variation might be due to the diverse risk factors as the practice of antibiotic use, increasing trend of prescribing third generation cephalosporins and type of the patients in the health care settings [31,32].

Overall prevalence of carbapenem resistance *Enterobacteriaceae* (CRE) was found to be 18.9%, which is lesser than the earlier reported observation from a tertiary care centre in Mumbai, 26.0% of the isolates were CRE [33]. However, the rate is higher compared to a similar study from parts of North and South India [34, 35]. Trend of carbapenem resistance was 28.2% among the *E. coli* isolates, which is higher than prevalence rate of 13.8% observed among the *Klebsiella* spp., contrasting the finding of previous study from Tamil Nadu [34]. These variations might be due to the selection bias or uneven distribution of the samples, sample size and the sample collection sites, included in the study.

## Conclusion

In conclusion, high prevalence of multidrug resistant, ESBL producer and carbapenem resistant *Enterobacteriaceae* isolates in the region is a menace, due to the risk of rooting out of the ultimate antimicrobials available. This emphasizes the urgent need of a comprehensive system to monitor and control the rapid spread of these super-bugs in this region.

## Acknowledgements

Our sincere thanks to Department of Biotechnology, Govt. of India for sanctioning the DBT-NER Twinning project (File No.BT/PR16669/NER/95/239/2015) and the Department of Life Sciences, Dibrugarh University for the facilities used during the research. We are thankful to Assam Medical College and Hospital authority especially the staff members of Department of Microbiology, for their kind support in sample collection process. The generous contributions of all the patients and the staff members of concerned hospitals in the project are also highly acknowledged.

## References

1. Nordmann P, Dortet L, Poirel L. Carbapenem resistance in in Enterobacteriaceae: here is the storm! Trends Mol Med. 2012; 18: 263–72.

2. Wang X, Chen G, Wu X, Wang L, Cai J, Chan EW, et al. Increased prevalence of carbapenem resistant *Enterobacteriaceae* in hospital setting due to cross- species transmission of blaNDM-1 element and clonal spread of progenitor resistant strains. Front Microbiol. 2015; 6: 595.

3. Nordmann P, Naas T, Poirel L. Global spread of Carbapenemase producing Enterobacteriaceae. Emerg Infect Dis. 2011; 17: 1791–8.

4. Moquet O, Bouchiat C, Kinana A, Seck A, et al. Class D OXA-48 carbapenemase in multidrug-resistant enterobacteria, Senegal. Emerg Infect Dis. 2011; 17: 143–4.

5. Roy, S., Viswanathan, R., Singh, A.K., Das, P., Basu, S. Sepsis in neonates due to imipenem resistant *Klebsiella pneumoniae* producing NDM-1 in India. J. Antimicrob. Chemother. 2011; doi: 10.1093/jac/dkr068

6. Bae IK, Kang HK, Jang IH, Lee W, Kim K, Kim Jo, et al. Detection of carbapenemases in clinical *Enterobacteriaceae* isolates using the VITEK AST-N202 card. Infect Chemother. 2015; 47: 167–74.

7. Djahmi N, Dunyach-Remy C, Pantel A, Dekhil M, Sotto A, Lavigne JP. Epidemiology of carbapenemase-producing *Enterobacteriaceae* and *Acinetobacter baumannii* in Mediterranean countries. Biomed Res Int. 2014; 305784.

8. November 2015 Update-CRE Toolkit, Facility Guidance for Control of Carbapenem-Resistant Enterobcateriaceae (CRE), 2015. https://www.cdc.gov/hai/organisms/cre/cre-toolkit/index.html

9. Mate PH, Devi KS, Devi KM, Damrolein S, Devi N.L, Devi P.P. Prevalence of Carbapenem Resistance among Gram-Negative bacteria in a Tertiary care Hospital in North East India. IOSR-JDMS. 2014; 13: 56–60.

10. Clinical and Laboratory Standards Institute (CLSI). Performance Standards for Antimicrobial Susceptibility Testing. USA: CLSI: M100-S25; 2015. Wayne, PA.

11. Shashwati N., Kiran T and Dhanvijay A.G. Study of extended spectrum β-lactamase producing Enterobacteriaceae and antibiotic coresistance in a teriary care teaching hospital. J Nat Sci Biol Med. 2014; 5: 30–5

12. Metri Basavaraj C, Jyothi P, Peerapur BasavarajV. The prevalence of ESBL among Enterobacteriaceae in a tertiary care hospital of North Karnataka, India. J Clin Diagn Res. 2011; 5: 470–5.

13. Rudresh SM, Nagarathnamma T. Extended spectrum β-lactamase producing Enterobacteriaceae and antibiotic co-resistance. Indian J Med Res. 2011; 133: 116–8.

14. Bradford PA. Extended-Spectrum β-Lactamases in the 21st Century: characterization, epidemiology, and detection of this important resistance threat. Clin Microbiol Rev. 2011; 14: 933–51.

15. Smet A, Martel A, Persoons D, Dewulf J, Heyndrickx M, Herman L, et al. Broad-spectrum β-lactamases among Enterobacteriaceae of animal origin: molecular aspects, mobility and impact on public health. FEMS Microbiol Rev. 2010; 34: 295–316.

16. Bush K, Fisher J. Epidemiological expansion, structural studies, and clinical challenges of new β-lactamases from gram-neative bacteria. Annu Rev Microbiol. 2011; 65: 455–78.

17. Dalela G. Prevalence of Extended spectrum beta lactamases (ESBL) producers among Gram Negative Bacilli from various clinical isolates in a tertiary care hospital at Jhalawar, Rajasthan, India. J Clin Diagn Res. 2012; 6: 182.

18. Teklu DS, Negeri AA, Legese MH, Bedada TL, Woldemariam HK, Tullu KD. Extended- spectrum beta-lactamase production and multi-drug resistance among Enterobacteriaceae isolated in Addis Ababa, Ethiopia. Antimicrob Resist Infect Control. 2019; 8: doi: 10.1186/s13756-019-0488-4

19. Thakur S, Pokhrel N, Sharma M. Prevalence of multidrug resistant enterobacteriaceae and extended spectrum β lactamase producing Escherichia coli in urinary tract infection. Res J Pharm Biol Chem Sci. 2013; 4: 1615–24.

20. Parajuli NP, Maharjan P, Parajuli H, et al. High rates of multidrug resistance among uropathogenic Escherichia coli in children and analyses of ESBL producers from Nepal. Antimicrob Resist Infect Control. 2017; 6: 9.

21. Nairouch YR, Mahafzah AM, Shehabi AA. Molecular Characterization of Multidrug Resistant Uropathogenic *E.coli* Isolates from Jordanian Patients. The Open Microbiol J. 2018; 12: 1–7.

22. Kulkarni SR, Peerapur BV, Sailesh KS. Isolation and antibiotic susceptibility pattern of Escherichia coli from urinary tract infections in a tertiary care hospital of North Eastern Karnataka. J Nat Sc Biol Med. 2017; 8: 176–80.

23. Eshetie S, Unakal C, Gelaw A, Ayelign B, Endris M, Moges F. Multidrug resistant and carbapenemase producing Enterobacteriaceae among patients with urinary tract infection at referral Hospital, Northwest Ethiopia. Antimicrob Resist and Infect Cont. 2015; 4: 12

24. Ahmed AJA, Haneen MRJA. Phenotypic and molecular characterization of multidrug resistant Klebsiella pneumoniae isolated from different clinical sources in Al-Najaf province-Iraq. Pak. J. Biol. Sci. 2017; 20: 217–32.

25. Metri BC, Jyothi P, Peerapur BV. The Prevalence of ESBL among Enterobacteriaceae in a Tertiary Care Hospital of North Karnataka, India. J Clin Diagn Res. 2011; 5: 470.

26. Bhattacharjee A, Sen MR, Prakash P, Gaur A, Anupurba S. Increased prevalence of extended-spectrum β lactamase producers in neonatal septicaemic cases at a tertiary referral hospital. Indian J Med Microbiol. 2008; 26: 356–60.

27. Das N, Borthakur AK. Antibiotic coresistance among extended-spectrum beta lactamase-producing urinary isolates in a tertiary medical center: A prospective study. Chron Young Sci. 2012; 1: 53–6.

28. Ravikant, Kumar P, et al. Prevalence and identification of extended spectrum β-lactamases (ESBL) in Escherichia coli isolated from a tertiary care hospital in North-East India. Indian J Exper Biol. 2016; 54: 108–14.

29. Izadi N, Nasab MN, Mood EH, Meshkat Z. Prevalence of TEM and SHV Genes in Clinical Isolates of Klebsiella Pneumonia From Mashhad, North- East Iran. Iranian J of Path. 2014; 9: 199 – 205.

30. Sharma M, Pathak S, Srivastava P. Prevalence and antibiogram of Extended Spectrum β-Lactamase (ESBL) producing Gram negative bacilli and further molecular characterization of ESBL producing *Escherichia coli* and *Klebsiella* spp. J Clin Diagn Res. 2013; 7: 2173–7.

31. Subha A, Ananthan S. Extended-Spectrum β-lactamase (ESBL) mediated resistance to third generation cephalosporins among Klebsiella pneumoniae in Chennai. Indian J Med Microbiol. 2002; 20: 92–5.

32. Jain A, Roy I, Gupta MK, Kumar M, Agarwal SK. Prevalence of extended-spectrum β-lactamase-producing Gram-negative bacteria in septicemic neonates in a tertiary care hospital. J Med Microbiol. 2003; 52: 421–5.

33. Nair PK, Vaz MS. Prevalence of Carbapenem resistant Enterobacteriaceae from tertiary care hospital. J Microbiol Infect Dis. 2013; 3: 201–10.

34. Sekar R, Srivani S, Amudhan M, Mythreyee M. Carbapenem resistance in a rural part of southern India: *Escherichia coli* versus *Klebsiella* spp. Indian J Med Res. 2016; 144: 781–3.

35. Datta P, Gupta V, Garg S, Chander J. Phenotypic method for differentiation of carbapenemases in Enterobacteriaceae: Study from north India. Indian J Pathol Microbiol. 2012; 55: 357–60.

